# Guidelines for obtaining an absolute blood flow index with laser speckle contrast imaging

**DOI:** 10.1101/2021.04.02.438198

**Authors:** Smrithi Sunil, Sharvari Zilpelwar, David A. Boas, Dmitry D. Postnov

## Abstract

Laser speckle contrast imaging (LSCI) is a technique broadly applied in research and clinical settings for full-field characterization of tissue perfusion. It is based on the analysis of speckle pattern contrast, which can be theoretically related to the decorrelation time - a quantitative measure of dynamics. A direct contrast to decorrelation time conversion, however, requires prior knowledge of specific parameters of the optical system and scattering media and thus is often impractical. For this reason, and because of the nature of some of the most common applications, LSCI is historically used to measure *relative* blood flow change. Over time, the belief that the absolute blood flow index measured with LSCI is not a reliable metric and thus should not be used has become more widespread. This belief has resulted from the use of LSCI to compare perfusion in different animal models and to obtain longitudinal blood flow index observations without proper consideration given to the stability of the measurement. Here, we aim to clarify the issues that give rise to variability in the repeatability of the quantitative blood flow index and to present guidelines on how to make robust absolute blood flow index measurements with conventional single-exposure LSCI. We also explain how to calibrate contrast to compare measurements from different systems and show examples of applications that are enabled by high repeatability.

## Introduction

Laser speckle contrast imaging (LSCI) provides a rapid wide field qualitative characterization of the motion of light scattering particles. Over the past 20 years, it has become widely used as an imaging tool to measure blood flow in the brain^1–6^, skin^7–9^, retina^10–12^ and other organs. LSCI has had a broad impact on fundamental research, for instance enabling the connection between migraine aura and headache^2^, and has a rising number of clinical applications^12,13^ thanks to the approval of LSCI imaging systems by the US Food and Drug Administration.

The technique is based on quantifying the speckle pattern blurring, caused by the motion of scattering particles, in terms of speckle contrast (K)^14^. The speckle contrast is a function of the exposure time (T) and the intensity autocorrelation function *g*_2_(*τ*). Their relationship is given by^15,16^:

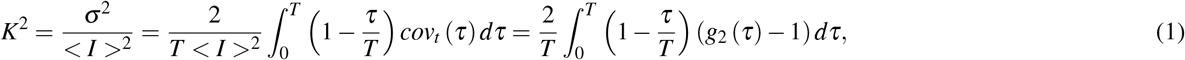

where *K* is the contrast, < *I* > and *σ* are the mean and standard deviation of the intensity respectively, *T* is the exposure time, *g*_2_(*τ*) is the intensity autocorrelation function, and *τ* is the time lag. The intensity autocorrelation function, in turn, is related to the field correlation function and thus to the decorrelation time *τ*_*c*_, a quantitative measure of the particles dynamics^14,16^. For spatial statistics in the presence of static scattering, the relation is commonly described as^16–18^:

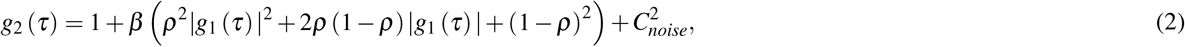

where the *β* parameter describes the degree of coherence^19–21^, *ρ* is the fraction of static scattering events^17,18,21–23^, and *g*_1_(*τ*) is the temporal field auto-correlation function^14^. Finally, 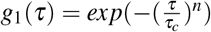, where *n* takes discrete values of 0.5, 1, or 2, depending on the form of the particle motion and the scattering regime^18,24^, and *τ*_*c*_ is the decorrelation time, which is inversely proportional to particles speed and thus to perfusion^16,25^. To allow real-time contrast estimates and to ease the adoption of the technique, the contrast to perfusion relation is generally expressed in terms of the blood flow index^11,20,26–30^ (BFI):

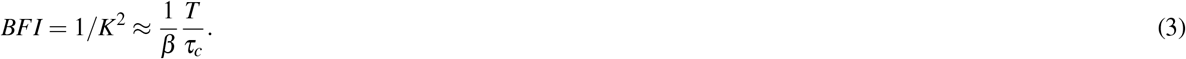

It is important to note that the parameters *ρ, n*, and *τ*_*c*_ describe the properties of the medium^17,18^, while *β* is the only parameter that depends on the imaging system characteristics^15,19,20^ and is not explicitly set by the user. To exclude system-dependent effects, the relative blood flow index (rBFI) is defined:

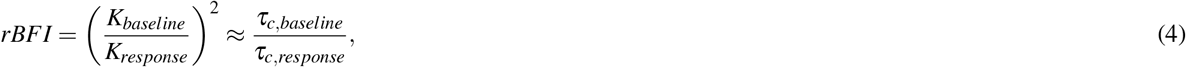

where baseline and response indicate either temporal (before vs after stimulus) or spatial (affected tissue vs non-affected tissue). As a system-independent quantitative metric, rBFI measured with LSCI has found broad applications in studying blood flow changes in various applications. These include, but are not limited to stroke experiments^3,6,28,29^, sensory stimulation^4,31^, perfusion response to drugs^11,32^, and vasoreactivity research^11,27^.

However, the use of absolute values of BFI for quantitative studies of flow has not evolved due to the BFI dependence on system parameters. As a result, LSCI has found a limited number of applications in studies where there is no baseline data or where it is challenging to avoid changes in the system configuration between experimental sessions. Such studies include long-term blood flow monitoring (i.e. stroke recovery, ageing), absolute perfusion comparison in different subjects or animal models, and studies in which animal has to be moved during the experiment (i.e. traumatic brain injury).

In this work, we aim to demonstrate that, when the system is appropriately configured, the absolute speckle contrast and resultant BFI are sufficiently robust for quantitative comparisons to be performed. It is important to note that we address the problem of quantitative BFI measurements and not the problem of relating BFI to the absolute perfusion values, which requires knowledge of the scattering regime and particle motion type in the media^18,25^. We demonstrate that measurement repeatability can be increased by adjusting polarization, speckle size, and the amount of light. We also show how contrast can be calibrated to enable comparisons between different settings and even different imaging systems. We conclude the results with two examples of applications where robust absolute blood flow index measurements, enabled by following our guidelines, provide important physiological insights.

## Results

### Repeatability of speckle contrast measurements

Key components and parameters of an LSCI system are shown in Fig.1. Some of them, such as sensor parameters (exposure time, frame rate), are easy to reproduce between studies and, in most cases, are well described. Others, such as optical parameters and the light source position relative to the sample, are more difficult to quantify and often vary even within the same set of experiments. The key characteristics that affect LSCI signal quality are: (i) exposure time *T* with respect to *τ*_*c*_, (ii) illumination intensity *I* with respect to the camera dynamic range and noise, and (iii) system parameters that define parameter *β*. The latter includes light source coherence and stability^20^, speckle to pixel size ratio^19^, and detected light polarization^19,33^. The *β* takes values from 0 to 1, where 1 corresponds to ideal conditions, and *β*<1 reflects the loss of correlation related to the listed factors. From eq. 2 it can be seen that increasing *β* is a key to improving the robustness and the signal-to-noise ratio of LSCI. Furthermore, keeping *β* constant between recordings is critical for the repeatability of LSCI measurements.

**Figure 1.**
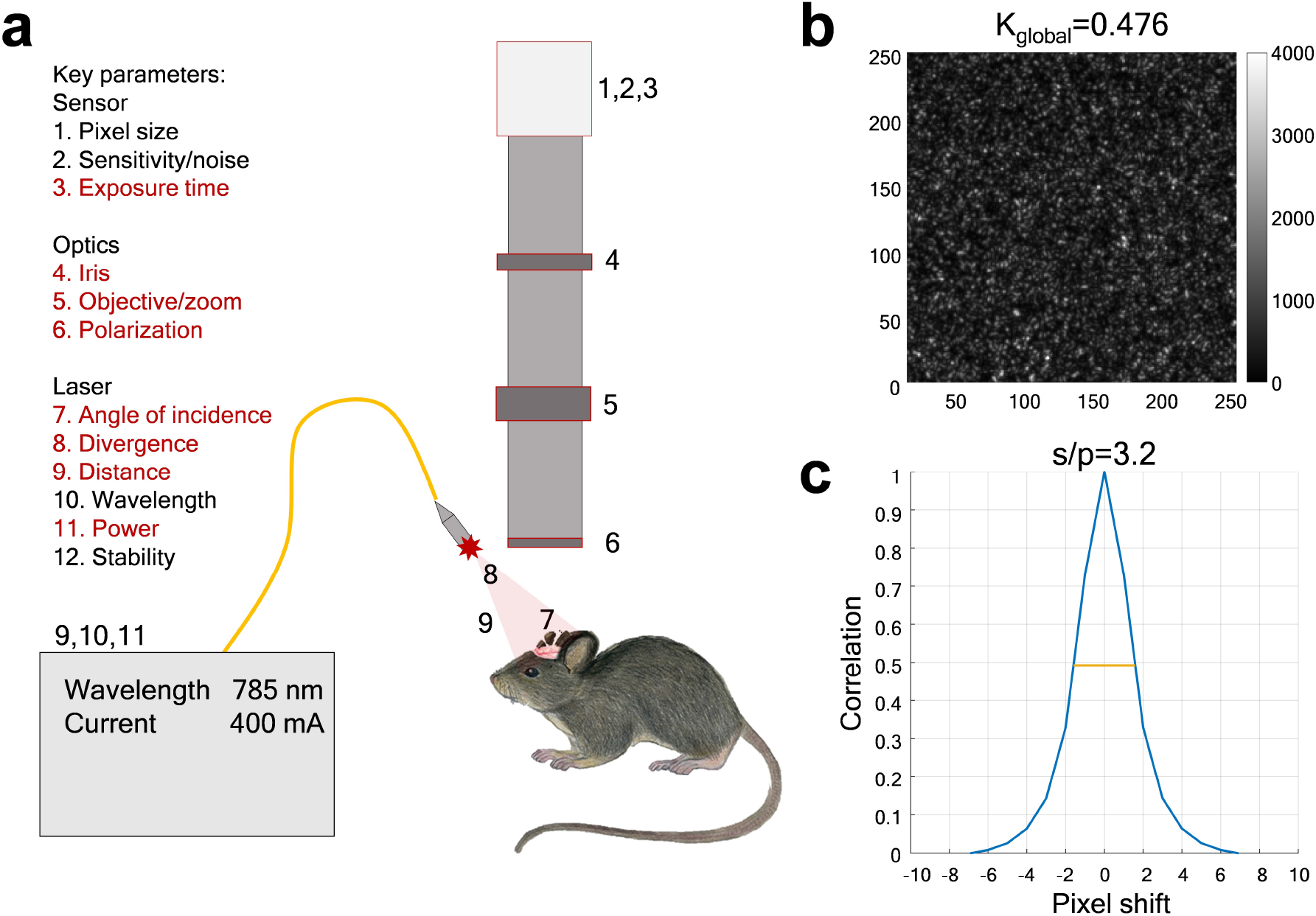
Key components and parameters of a typical LSCI system. (a) List of the LSCI system components. Parameters highlighted in red are often modified between recordings even within the same system. (b), (c) Speckle pattern characteristics that vary with the LSCI system parameters: average intensity, global contrast, and speckle to pixel size relation *s/p* (as determined from the spatial cross-correlation of the speckle pattern).

To quantify repeatability we introduce two metrics - *D*_*m*_ and *D*_*sd*_. *D*_*m*_ is a metric of the standard deviation of contrast between recording sessions relative to the mean. *D*_*sd*_ is a metric of the standard deviation of contrast between recording sessions relative to the average standard deviation of the contrast within recording sessions.

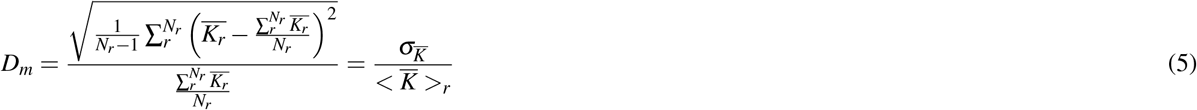

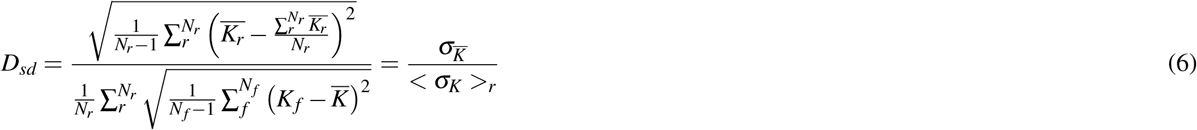

where *K*_*f*_ is the spatial contrast calculated for the frame *f, N*_*f*_ is the total number of frames, 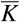 is the average spatial contrast across all frames within a single recording, *σ*_*K*_ is the standard deviation of the spatial contrast for all frames in a single recording, and 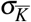 is the standard deviation of the average spatial contrast between recordings. *N*_*r*_ is the total number of recordings, 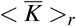 is the contrast averaged across all frames in all recordings, and < *σ*_*K*_ >_*r*_ is the mean of the standard deviation within recordings.

Example contrast measurements and their repeatability are shown in Fig.2. The system parameters are: the speckle to pixel size ratio (s/p) of 3.2; the detected light is polarized; a VHG stabilized laser diode with a coherence length ≈ 2mm is used as the light source. Parameters remain constant between recordings, except for natural changes in the position of the sample as two animals were alternated. Under these conditions, the repeatability is high with the contrast variation between recordings being less than 6% of the mean (*D*_*m*_<0.06, see Fig.2, c), and less than 25% of the variation within the recordings (*D*_*sd*_<0.25, see Fig.2, d) caused by measurement noise and physiological fluctuations (e.g. breathing, cardiac activity).

**Figure 2.**
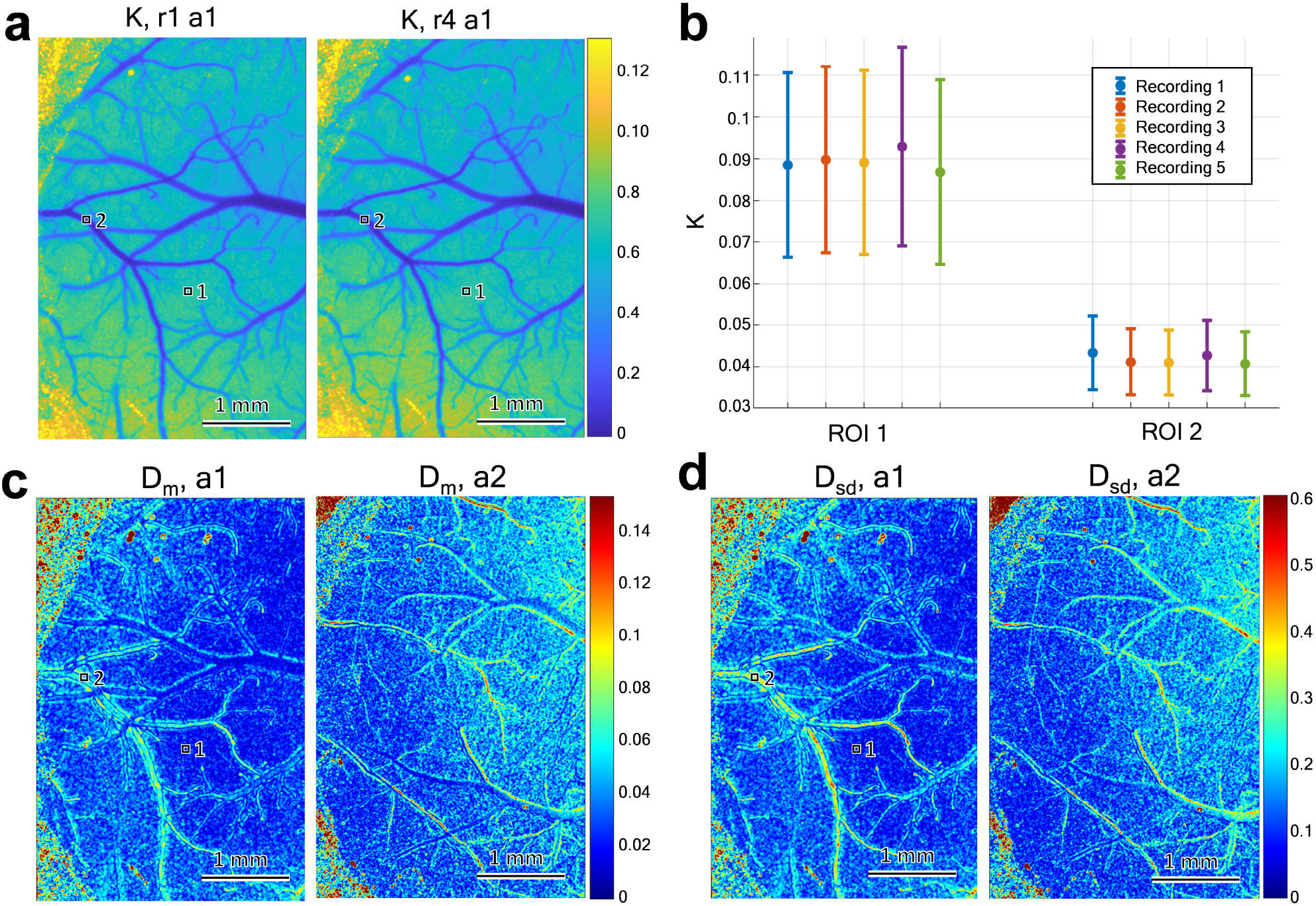
Repeatability in the speckle contrast measurements. (a) Spatial contrast map measured from the same animal after repositioning. (b) Average and standard deviation of the speckle contrast values in selected ROIs for five different recordings from the same animal. (c) and (d) *D*_*m*_ and *D*_*sd*_ repeatability metrics for recordings in both animals. The variation between recordings is minor compared to the average contrast level and the average variation within given recordings - in majority of the pixels *D*_*m*_<0.06 and *D*_*sd*_<0.25.

To further explore the repeatability, we have performed additional experiments, where the system parameters were varied in addition to alternating animals and adjusting their position. The s/p was adjusted to take values 2.5,2.8,3.7 or 5.8, and the polarization of the detected light was set to either “no polarization” (by removing the polarizing filter) or linearly polarized to minimize the reflected light. We took six recordings (*N*_*r*_ = 6) for each animal with each of 8 parameter sets. As exposure time was kept constant and s/p was changed by opening or closing the iris, the average intensity < *I* > varied between recordings. Fig. 3,(a) shows the pixel average *D*_*m*_ and *D*_*sd*_ for all recordings, including the ones demonstrated in Fig. 2. One can see that even the least robust settings (s/p=2.5 with non-polarized detection) result in *D*_*m*_ < 0.14 and *D*_*sd*_ < 0.45, which can be considered acceptable for studies where large changes in perfusion are expected. The repeatability improves with larger s/p ratios and polarized detection, reaching *D*_*m*_ < 0.03 and *D*_*sd*_ < 0.13. Recordings with high repeatability tend to share the same characteristics: detected light is polarized, the average intensity variation is low, and the s/p value is typically above 2.8. Fig 3,(b) and (c) show <u>ho</u>w the repeatability metrics are affected by the size of the neigh<u>b</u>ourhood used to calculate the contrast. Average contrast 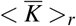 rapidly rises with the increase of the neighbourhood side 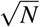 (where N is the total number of pixels in the neighbourhood) and reaches 80% of possible maximum when 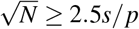, which agrees well with computational studies^33^. The *D*_*m*_ rapidly decreases (improvement of repeatability) until the maximum 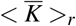 is reached, which is followed by a slower decrease caused by changes in 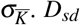. *D*_*sd*_, on the other hand, is rising, reflecting the reduction of noise effects and decreased average variation within recording < *σ*_*K*_ >_*r*_.

**Figure 3.**
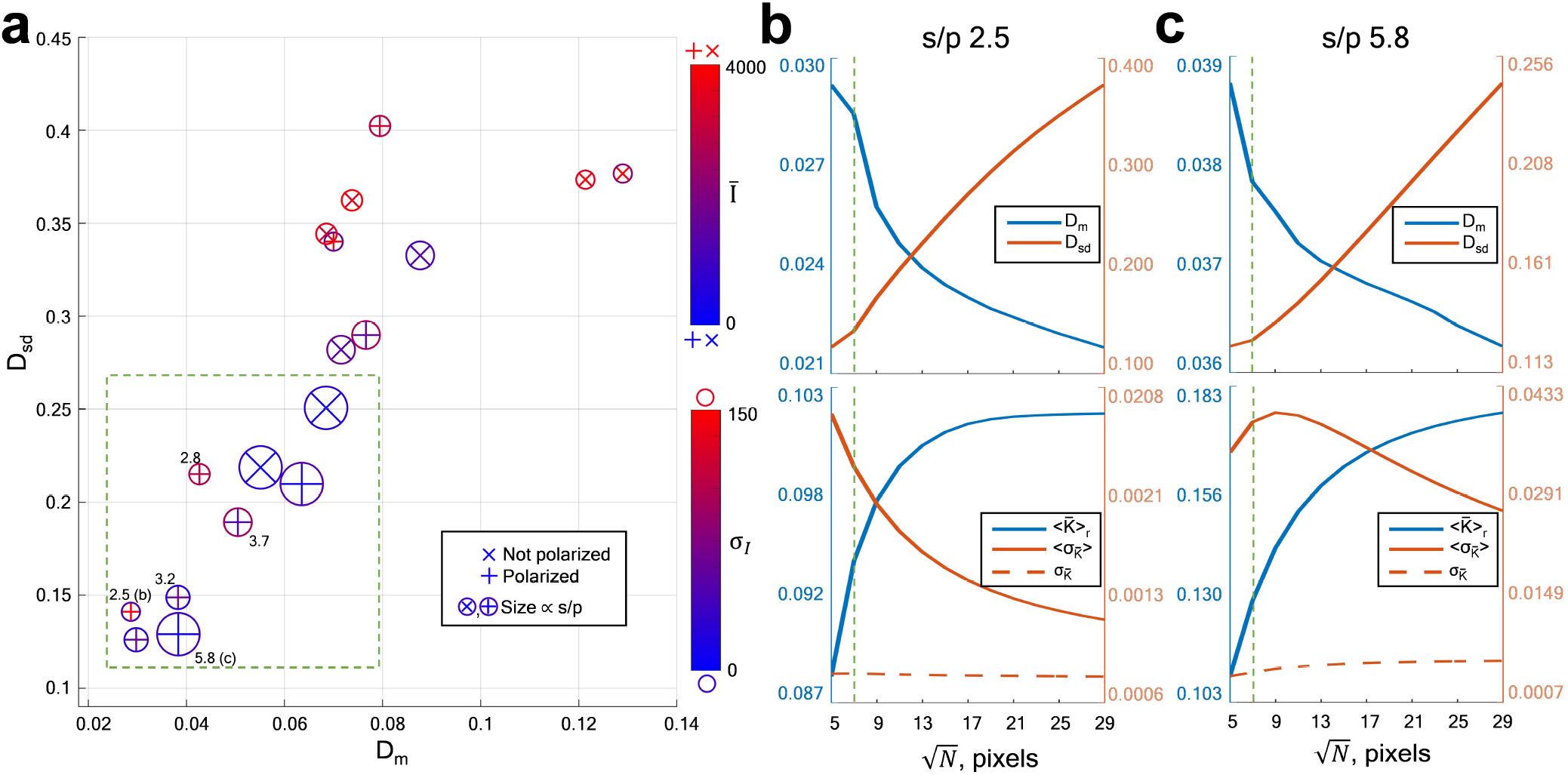
(a) Pixel average repeatability metrics for recordings taken with varied system parameters. Circle size is proportional to the *s/p* value and corresponds to 2.5, 2.8, 3.2, 3.7, and 5.8. The symbol inside the circle denotes data taken with the polarizing filter (’+’) and without it (’x’), with the symbol colour indicating the average intensity. The colour of the circle outline indicates the standard deviation of the average intensity between recordings. Green dashed square highlights 9 out of 18 recordings with the highest repeatability. (b),(c) Pixel average repeatability metrics for the spatial contrast calculated using neighbourhoods of different size. Recordings with high repeatability and speckle to pixel ratios 2.5 (b) and 5.8 (c) were analyzed. Corresponding system settings are indicated in (a), bottom left corner. Green dashed line highlights the commonly used 7×7 (N=49) pixels neighbourhood.

### Cross-parameter and cross-system calibration

An estimate of *β* is required to calibrate and compare BFI (or speckle contrast) measured with different parameters or imaging systems. It can be obtained by imaging a static scattering phantom (see Fig.1,b) and calculating the global contrast^20^. According to eq.1,2, the speckle contrast measured from a purely static phantom (*τ*_*c*_ = 0) will be given by:

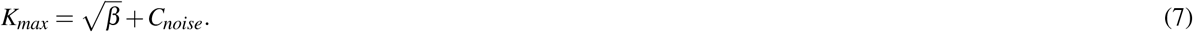

*K*_*max*_ is the maximum speckle contrast that can be measured with the given system parameters, as any dynamics in the scattering particles will only reduce the contrast. Given *K*_*max*_, we can now calibrate subsequent measurements as:

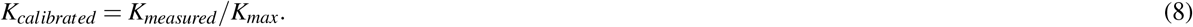

We applied the calibration procedure and calculated pixel average *D*_*m*_ for all possible pairs of recordings taken with the settings shown in Fig.3. Results showing the impact of calibration are presented in Fig.4. As expected, changing the speckle size or polarization often leads to high contrast variation between the recordings (*D*_*m*_ reaches 0.3, Fig.4, (a) left panels), which will prevent perfusion comparison between the datasets. The difference is particularly noticeable when comparing two different polarization states, or comparing the smallest speckle size (*s/p* = 2.5) with larger speckle sizes (*s/p* = 3.7 and 5.8). The difference between recordings taken with *s/p* = 3.7 and *s/p* = 5.8 is not as noticeable (*D*_*m*_<0.1), since effects of speckle to pixel size ratio on the average contrast becomes negligible after *s/p* reaches 3^19^. After the calibration (Fig.4, (a) right panels)), the *D*_*m*_ generally reduced to below 0.1 within the same polarization - an example is shown in Fig. 4, b. Calibration becomes less effective when applied to measurements taken with different polarizations - repeatability has improved in 28 out of 32 cases (Fig.4,c), but worsened in other 4 (all observed in animal 2, Fig.4,e). Importantly, calibration allows comparing the data collected with two different systems as long as the *K*_*max*_ estimate is available for both systems (Fig.4,d).

**Figure 4.**
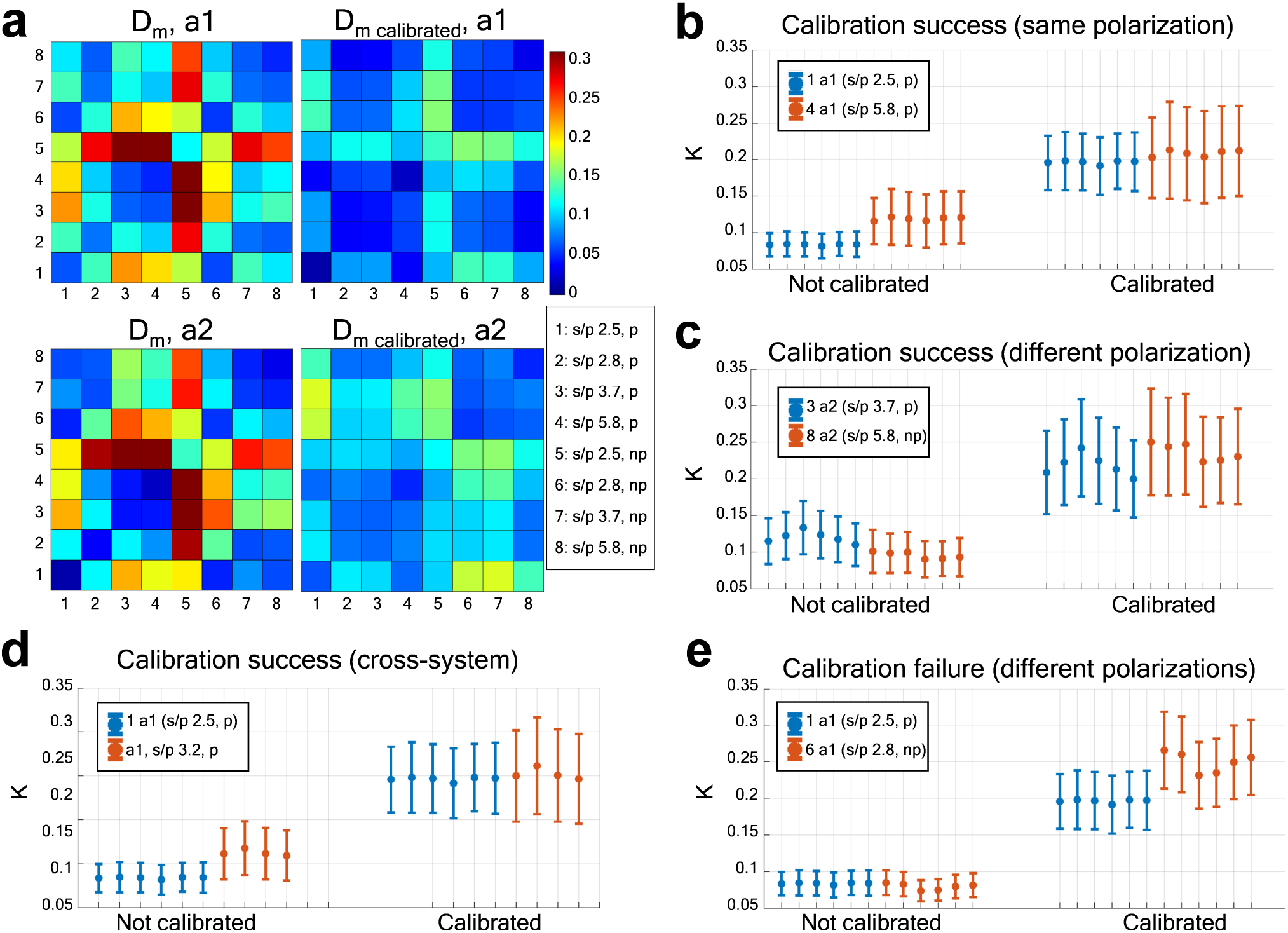
Contrast calibration results. (a) The pixel average *D*_*m*_ and *D*_*m calibrated*_ for both animals, calculated with the data merged from recordings taken with different settings. Calibration improves the repeatability, especially when applied to the data collected with the same polarization type. (b)-(e) Examples of the calibration displayed as the average and standard deviation of the contrast over the field of view. (b) Calibration of the data collected with the same polarization type; (c) successful calibration with different polarization types; (d) successful calibration between different imaging systems (same polarization type); (e) failed calibration for the data collected with different polarization types.

### Potential applications

In Fig.5 we present examples of both long term and short term recordings where repeatable LSCI is necessary or beneficial. Fig.5 (a),(c),(e) show long term monitoring of cerebral blood flow in the animal model of ischemic stroke^3^, where continuous imaging is challenging due to the long, over few days, observation time. Fig.5 (b),(d),(f) show monitoring of cerebral blood flow during a traumatic brain injury experiment, where continuous imaging is not possible as the experimental design requires moving the animal between different setups. In the stroke experiments the control ROI returned to the baseline perfusion level at 24 hours after stroke and remained at that same level at the 72 hr time-point, while the ischemic core remained at *rBFI* = 0.5. Without ensuring repeatable LSCI, one can only compare between ROIs within a single time point, which might result in the misinterpretation of the results as both regions are affected to a different degree depending on the time passed after the stroke. With repeatable LSCI, more robust analysis is possible by comparing each time point to the pre-stroke blood flow index. Similarly, in the traumatic brain injury experiment, repeatable LSCI allows evaluation of the blood flow reduction by comparing it to the baseline instead of the control hemisphere, where the blood flow also changes over time.

**Figure 5.**
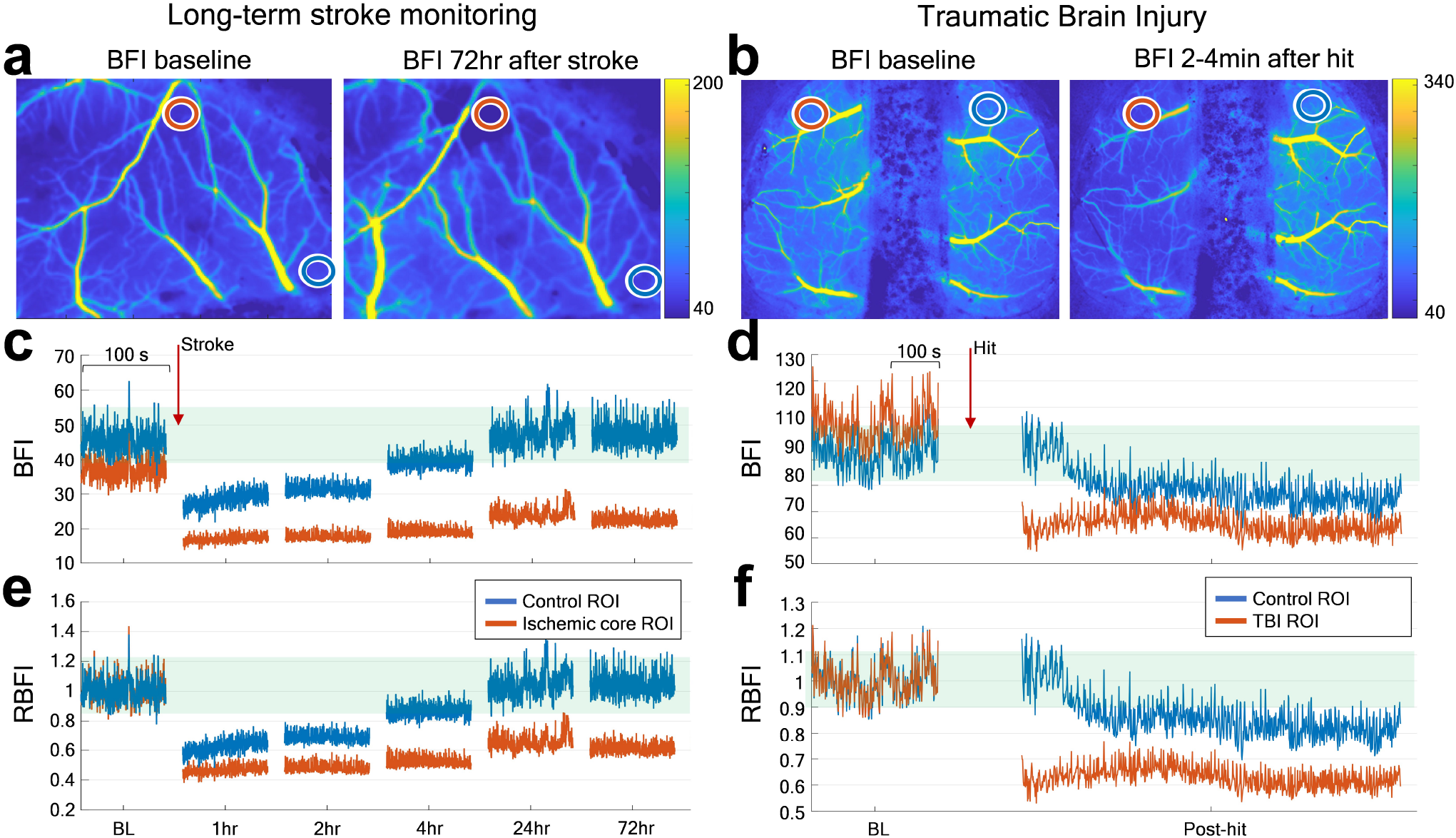
Application examples, where continuous imaging is challenging due to requirements of the experimental design. (a),(c),(e) Long term monitoring of cerebral blood flow in the animal model of ischemic stroke. (b),(d),(f) Monitoring of cerebral blood flow during a traumatic brain injury experiment. Orange lines and ROIs correspond to the ischemic core (in the stroke experiment) or the hemisphere that was hit (in the traumatic brain injury experiment). Blue lines and ROIs correspond to the control regions, which are not affected directly.

## Discussion

We have demonstrated that the absolute values of speckle contrast and the corresponding BFI collected with an optimally configured LSCI system are sufficiently robust and repeatable for quantitative comparison. We have also shown how to calibrate the contrast to allow comparison of BFI measurements taken with different imaging systems (i.e. by different research groups). To achieve the optimal configuration one should follow the guidelines presented below.

### Guidelines

- Use a linear polarizing filter in the detection path. Adjust the filter rotation to minimize the specular reflections, which generally corresponds to the minimum average intensity observed during the filter rotation. Although the importance of polarizing the detected light in the LSCI is well known^16,19^, this step is often missing in the real-life applications.
- Use a highly coherent and stable light source. For example - volume holographic grating wavelength stabilized diodes coupled to a single-mode fibre, that have a long coherence length and have become broadly available over the past few years.
- Have a detection pupil with an adjustable diameter that will permit adjustment of the system f-number and thus speckle size. Theoretically, a value of 2 is sufficient for the speckle to pixel size ratio to minimally affect *β* ^19^. However, in practice, because of the pixels cross-talk^20,34^ and difficulties in precise estimation of the speckle size, we recommend adjusting the pupil diameter to reach a speckle to pixel size ratio *s/p*≳ 3. The *s/p* value can be calculated by analyzing spatial cross-correlation of a speckle pattern recorded from a static scattering sample (Matlab script is available at BU NPC GitHub^35^). The same data can be used to estimate *K*_*max*_.
- When calculating the spatial contrast use neighbourhood side 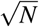 at least 2-3 times larger than the speckle to pixel size ratio (i.e. 7×7, *N*=49 for the *s/p*=3). Further increase in the neighbourhood size might be beneficial in the applications where spatial resolution loss is not of critical importance (i.e. skin imaging).
- Use an exposure time optimized according to the dynamics of interest: from ≈1 ms if measuring the blood flow index in arteries and veins with a diameter more than 50 micrometres, to ≈5 ms for capillaries and small vessels^36^ (e.g. imaging the skin or parenchymal regions of the brain).
- The average detected intensity over the field of view should be just below the middle of the sensor’s dynamic range (i.e. 110 for an 8 bit sensor). Lower values will result in a reduced ability to resolve speckle fluctuations, while higher average intensity might lead to the pixel saturation and clipping of the signal.
- Illuminate the sample as homogeneously as possible to ensure that the same signal to noise ratio is obtained over the entire field of view.
- If relevant, collect the data at a higher frame rate to allow contrast averaging in time and thus reduction of the shot noise effects. Compensate for the fixed pattern noise if it is noticeable. It increases the intensity of specific pixels, and thus impacts the relationship between calculated contrast and the perfusion.
- Collect the calibration data (speckle pattern image collected from a static scattering sample) and calculate *K*_*max*_ whenever the system parameters were or could be changed. Follow the same guidelines as described above during the calibration and modify the exposure time *T* to adjust the pixel intensity. Repeat the calibration in fixed intervals of time (i.e. once per week) during long term studies. Use the calibrated BFI 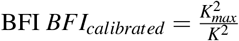 to evaluate perfusion.
- To make results reproducible by others, report the calibration parameter *K*_*max*_ and exposure time *T* along with the contrast or BFI measurements.

### Potential applications

Applications, where repeatable measurements with LSCI are critical, include perfusion comparison without stimulation and experiments where continuous monitoring is not possible. It will be particularly crucial for long term perfusion monitoring, such as in chronic stroke studies, where potential physiological changes compromise the use of control ROIs. Optimizing LSCI measurements will improve quality and open new possibilities for individual studies while publishing calibration data will allow BFI comparisons between different studies. Thus linking together the body of LSCI measurements performed in the world and allow the quantitative use of the LSCI blood flow index.

## Methods

Two separate LSCI systems were used. Both systems use the same model of the light source (volume holographic grating stabilized laser diode, 785 nm, LP785-SAV50, Thorlabs) to deliver coherent light to the sample and the same model of the CMOS camera (Basler acA2040-90umNIR, 2048×2048 pixels, 5.5 *µ*m pixel size) to record the backscattered light. The light was delivered to the camera through a 2x objective with NA= (TL2X-SAP, Thorlabs) in the first system, and a 5x objective with NA= 0.14 (Mitutoyo, Japan) in the second system. The speckle size of the first system was adjusted by altering the pupil diameter of an iris in the detection path to achieve an estimated speckle to pixel size ratios of ≈ 2.5,2.8,3.7 and 5.8. For the second system, the speckle to pixel size ratio was fixed to an estimated value of 3.2. A linear polarizing filter was either placed in front of the objective and adjusted to minimize the reflected light (polarized state), or removed from the system entirely (non-polarized state). The light source was fixed in the position that provided homogeneous illumination over the whole field of view and resulted in the optical power density that saturated 1% of pixels at the smallest speckle to pixel size ratio for the non-polarized system settings. As we did not change the optical power density between recordings, the average intensity is decreased for both the polarized condition and for larger speckle to pixels size ratios (smaller pupil diameter).

In all experiments, except for the application demonstration (stroke and traumatic brain injury), data were collected over 30 seconds per recording. The frame rate was 50 frames per second, and the exposure time was 5ms. The raw data was then processed into spatial contrast *K* by calculating standard deviation and mean of intensity over a neighbourhood of 7×7 pixels^16,33^ unless indicated otherwise. The contrast was then further processed to calculate maps of *D*_*m*_ and *D*_*sd*_ as well as *BFI* and *rBFI*, according to equations 3,4,5,6. All animal procedures were approved by the Boston University Institutional Animal Care and Use Committee and conducted following the Guide for the Care and Use of Laboratory Animals. A chronic cranial window surgery was performed on wildtype C57Bl6 mice (approximately 12-week-old, The Jackson Laboratory), as reported previously^3^. Briefly, a craniotomy was drilled over one hemisphere, keeping the dura intact. A half-skull shaped curved glass (modified from Crystal Skull, LabMaker, Germany) was inserted into the craniotomy, gently touching the brain.

The window was fixed with dental acrylic, along with an aluminium bar attached to the skull on the other hemisphere for head fixation during imaging. In the application experiment that involved traumatic brain injury, a half skull glass window was also installed in the second (control) hemisphere. In all experiments, except the application demonstration, animals were imaged under isoflurane anaesthesia (3% induction, 1-1.5% maintenance, in 1 L/min oxygen), by fixing the head to a custom-made stage. The animal was allowed to recover after the experiment. In the experiments that involved stroke and traumatic brain injury, after a 7-day surgery recovery period, the animals were trained for 10 days to remain awake and head-fixed under the imaging systems. The application experiments were performed in awake head-fixed mice.

The static scattering sample (white backside of the VRC2 Thorlabs laser viewing card) was imaged to get the calibration data. A single image was collected and processed to get the *K*_*max*_ value by calculating global contrast over 512×00512 pixels. The calibration data was collected for each speckle to pixel size ratio in the polarized state. To calibrate non-polarized data, *K*_*max*_ measured in the polarized state was multiplied by a theoretical coefficient of 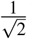 ^19,33^

## Acknowledgements

D.D. Postnov was supported by grant NNF17OC0025224 awarded by Novo Nordisk Foundation, Denmark. Support was also provided by NIH R01-MH111359, R01-EB021018, and R01-NS108472.

## Author contributions statement

S.S. and S.Z. prepared the animals, conducted the experiments, analyzed the results. D.A.B. conceived the study. D.D.P conceived the study, experiments, analyzed the results. All authors participated in writing and reviewing the manuscript.

## Additional information

## Data availability

The datasets generated during and/or analyzed during the current study are available from the corresponding author on reasonable request.

